# Evolutionary trade-offs between intergenerational and transgenerational fitness effects

**DOI:** 10.1101/2025.11.04.686475

**Authors:** Isaac Harris, Elizabeth M. L. Duxbury, Tracey Chapman, Simone Immler, Alexei A. Maklakov

## Abstract

Intergenerational and transgenerational fitness effects can shape evolutionary processes. Theoretically, however, intergenerational and transgenerational effects can trade-off with each other with profound consequences for evolutionary processes. Here we show that beneficial intergenerational effects that increase offspring fitness can result in detrimental transgenerational effects that decrease great-grand offspring fitness. We combined theoretical modelling and experimental approaches to investigate multigenerational fitness trade-offs induced by larval starvation in *Caenorhabditis elegans*. We demonstrate that larval starvation triggers a cascading effect: starved individuals suffered marked fitness losses, their direct offspring enjoyed fitness gains in both starvation and ad libitum environments, but great-grand-offspring paid fitness costs. Demographic simulation models revealed that genotypes exploiting this short-term intergenerational advantage outcompete rival genotypes despite the deferred transgenerational debt. Our findings demonstrate that adaptive intergenerational gains can be intrinsically linked to maladaptive transgenerational outcomes, challenging the assumption that transgenerational effects are inherently beneficial and highlighting the role of multigenerational trade-offs in evolution.

## Introduction

Intergenerational and transgenerational effects occur when the environmental experiences of parents or more distant ancestors influence the fitness of subsequent generations, potentially shaping evolutionary trajectories and ecological dynamics^1–6^. While intergenerational effects involve immediate parental-offspring interactions that typically enhance offspring fitness, transgenerational effects extend beyond direct offspring, influencing fitness in later descendants. By convention, intergenerational effects are present in the F_1_ and F_2_, while transgenerational effects occur in the F_3_ generation and onwards_7_. These effects have been commonly viewed as adaptive mechanisms allowing organisms to adjust their descendants’ phenotypes to anticipated environmental conditions. Several studies have described potentially adaptive intergenerational effects, such as decreased oxidative stress under parasite load in birds_8_, increased reproductive fitness during drought in plants_9_, and defensive head morphs under predator stress in *Daphnia*_10_. However, in contrast, empirical evidence for adaptive transgenerational effects remains limited_11–15_.

Theoretical models have described two adaptive evolutionary hypotheses of transgenerational effects. Anticipatory effects ‘pre-arm’ offspring, conferring fitness benefits to offspring experiencing the same stressful environment as their parents_16_, while heritable bet-hedging increases offspring trait variance and, therefore, population survivability, through trading an immediate decrease of arithmetic mean fitness for an increase in, long term, geometric mean fitness_17_. While theoretical models provide compelling explanations for the adaptive value of transgenerational effects, further empirical tests are needed to assess their generality and ecological significance in the face of putative evolutionary trade-offs_18_.

Inter- and transgenerational effects can transiently alter life-history traits. For instance, changes in parental diet can alter offspring lifespan and reproduction in *Drosophila*_19_ and survival in nematodes_20_. Because an organism’s resources in nature may often be limited, investment into a life-history trait may result in trade-offs with other traits. Such resource allocation trade-offs are a core component of life-history theory and can accompany phenotypic responses to environmental changes_21_. The fitness consequences of such trade-offs have been explored both within parental generations (P_0_)^22^ and their immediate offspring (F_1_)^23^. For instance, Burton and colleagues found offspring from nematodes exposed to osmotic stress exhibit intergenerational adaptations to oxidative stress but suffer a decrease in their pathogen response, giving evidence for an intergenerational trade-off between two phenotypically plastic traits_23_. Whilst existing work on multigenerational effects has focused on trade-offs between parents and their immediate offspring, the fitness consequences of long-term trade-offs remain largely unexplored. Such “multigenerational trade-offs” could arise when an adaptive effect in one generation of descendants (e.g. F_1_) imposes a fitness cost in a later generation (e.g. F_3_) or vice-versa. Despite their potential significance for understanding how populations react to stress in changing and fluctuating environments, direct empirical tests for such multigenerational trade-offs are scarce.

Here we used experimental and theoretical approaches to investigate intergenerational and transgenerational effects following larval starvation-induced developmental arrest in the nematode *Caenorhabditis elegans* (*figure 1*). Previous work has shown starvation during the first larval stage (L1) results in an intergenerational increase in offspring provisioning_24,25_ and a transgenerational increase in starvation survival_26_. By taking multiple fitness components (lifetime reproductive success, age-specific reproduction, rate-sensitive fitness and survival) across match-mismatched environments, we found that, independent of their environmental conditions, descendants of starved individuals exhibit increased intergenerational (F_1_) fitness but decreased transgenerational (F_3_) fitness.

**Figure 1:**
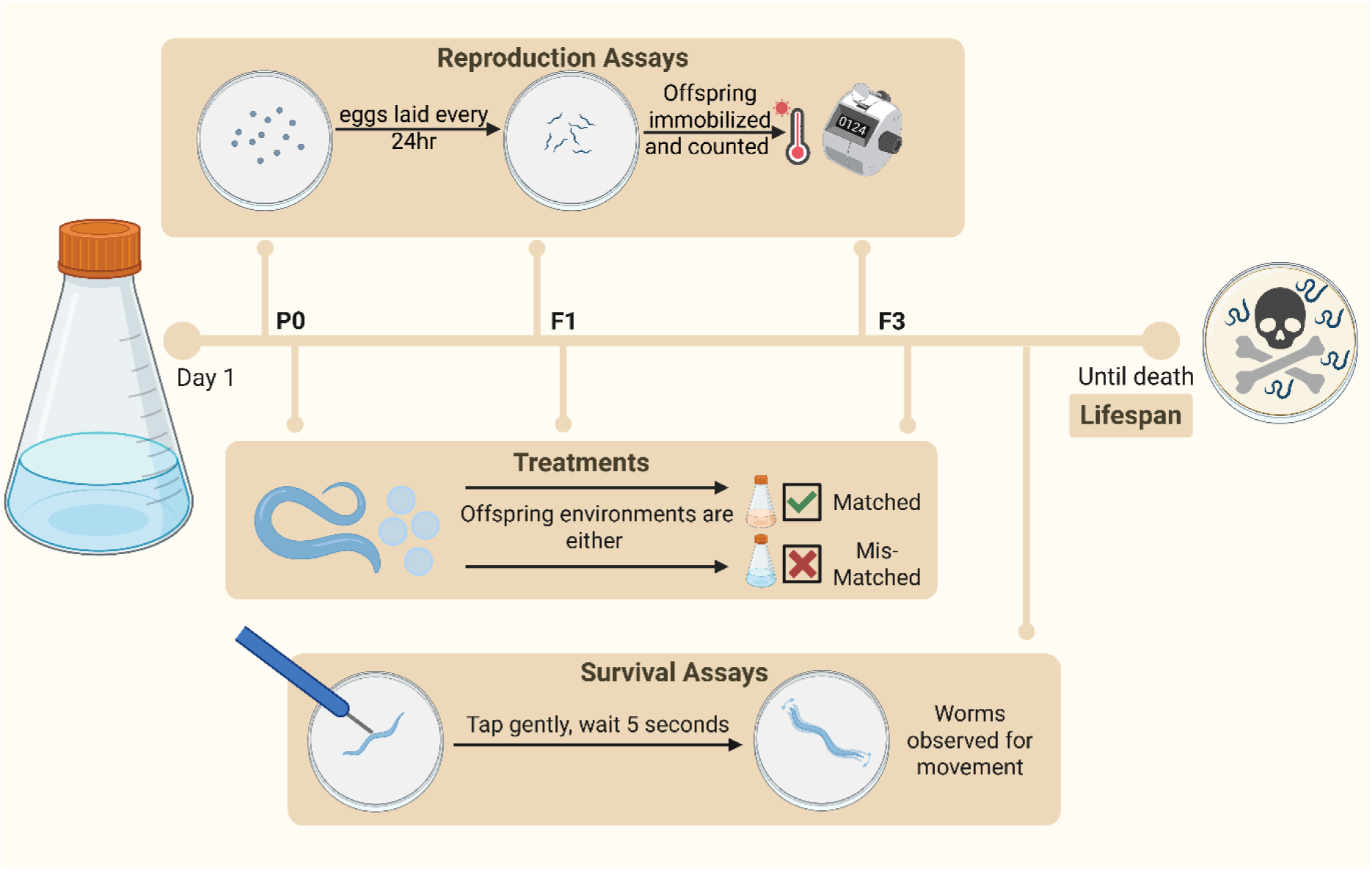
Experimental Design. We placed fertilized *Caenorhabditis elegans* eggs into either an environment with no food or *ad-libitum* food and allowed them to hatch. Worms which hatched in an environment with no food enter developmental arrest. We allowed worms to develop in *ad-libitum* conditions after 7 days of developmental arrest or after 24 hours in control conditions. F1 and F3 offspring of P0 worms were either put back into the same or the opposite environment to the P0 worms. F1 and F3 lines were run independently to each other and the F3 line was allowed to develop as normal through F1 and F2 generations. Fitness assays were taken in P0, F1 and F3 generations. For reproduction assays, worms were transferred to new plates every 24hrs, eggs were allowed to hatch and develop, and before the offspring reached reproductive age individuals were immobilized and each brood counted. Survival was assayed by gently touching worms each day and observing any movement. Worms unresponsive to touch were classed as dead. Figure was created with BioRender.com

Evolutionary simulation models of boom-and-bust population dynamics showed that maladaptive transgenerational effects can evolve as a cost of beneficial intergenerational effects. Together, our findings challenge the assumption that transgenerational effects are inherently beneficial and advance our understanding of how environmental stress can alter population dynamics in changing environments.

## Results

### P0

#### Age-Specific Reproduction

Reproductive timing was significantly affected by extended seven-day L1 larval starvation in the parental generation (*Anova type III: chi-squared χ2 = 33.39, p < 0.001*), as indicated further by the linear and quadratic interactions with age (*EMM wald z-test: z = 12.49, p < 0.001; z = -6.63, p < 0.001*). Individuals that experienced extended larval starvation had a delayed reproductive schedule compared to individuals raised in control conditions. Larval starvation resulted in a one-day delay in both peak reproduction and onset of reproduction (*Figure 2c*), with most individuals starting reproduction on day 2 and peaking on day 3. Moreover, peak reproduction was decreased in larval starvation treatments.

**Figure 2:**
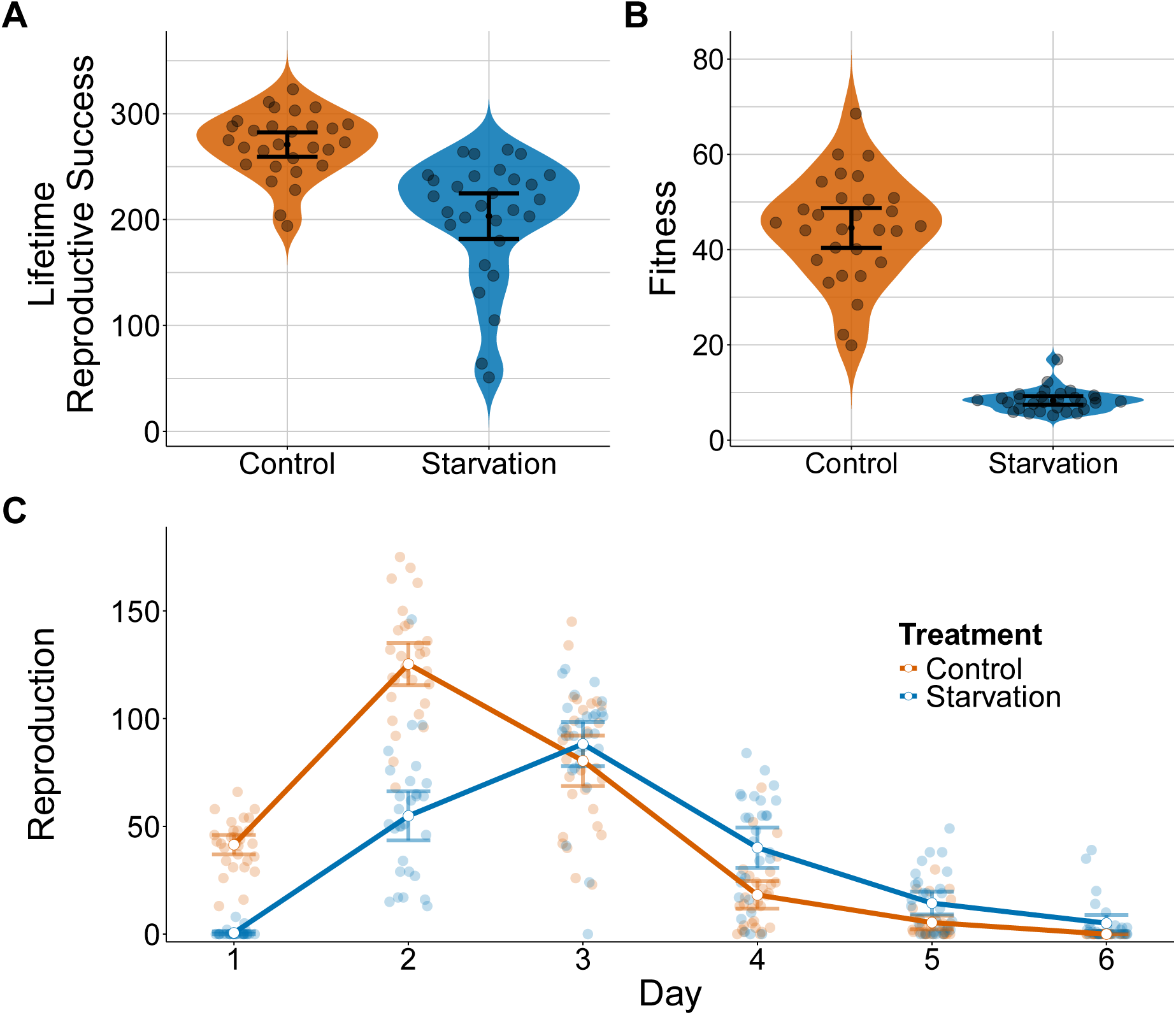
P0 Fitness Assays. P0 worms developed in either control (*ad-libitum*) or larval starvation conditions. **A&B)** The lifetime reproductive success and rate sensitive fitness (λ) respectively **C)** Plotted is daily reproduction, with line connecting the mean values for each treatment on each day. Each point represents one individual. All error bars represent 95% CIs.

#### Fitness

Similarly, lifetime reproductive success was significantly decreased in individuals that experienced extended larval starvation (*EMM wald z-test: z = -4.25, p < 0.001*) (*Figure 2A*). The delay in reaching peak reproduction along with an overall decrease in total reproduction resulted in a lower rate-sensitive fitness, *λind,* for individuals subjected to larval starvation (*EMM wald z-test: z = -24.13, p < 0.001*) (*Figure 2B*).

#### Survival

The survival of P_0_ individuals was significantly reduced by larval starvation (*EMM wald z-test: z = 2.31, p = 0.0209*) (*Figure S1*). Individuals who underwent larval starvation exhibited significant increases in risk of mortality (hazard), in comparison to individuals who developed in *ad libitum* conditions (*Log Odds =-0.61,95%: -1.12, - 0.092, p = 0.0207*).

### F1

#### Age-Specific Reproduction

Reproductive timing was significantly influenced by treatment (*Anova type III: χ2 = 8.19, p = 0.04*) (*Figure 3E*). Interactions with age were significant on both the linear and quadratic scale (*Anova type III: χ2 = 105.20, p < 0.001; χ2 = 46.23, p < 0.001*). Larval starvation in the F_1_ generation caused delayed peak reproduction regardless of parental treatment (**Offspring treatment (Parental treatment)**, EMM walds z test: **F_1_Control(P_0_Control) – F_1_Control(P_0_Starvation)**: z = 6.17, p <0.001, **F_1_Starvation(P_0_Control) – F_1_Starvation(P_0_Starvation)**: z = 6.79, p < 0.001). However, worms from starved parents reproduce more earlier than those from control parents regardless of their environment (EMM wald z-test: **F_1_Control(P_0_Control) – F_1_Starvation(P_0_Control)**: z: -13, p < 0.001, **F_1_Control(P_0_Starvation)-F_1_Starvation(P_0_Starvation)**: z: -11.84, p < 0.001)

**Figure 3:**
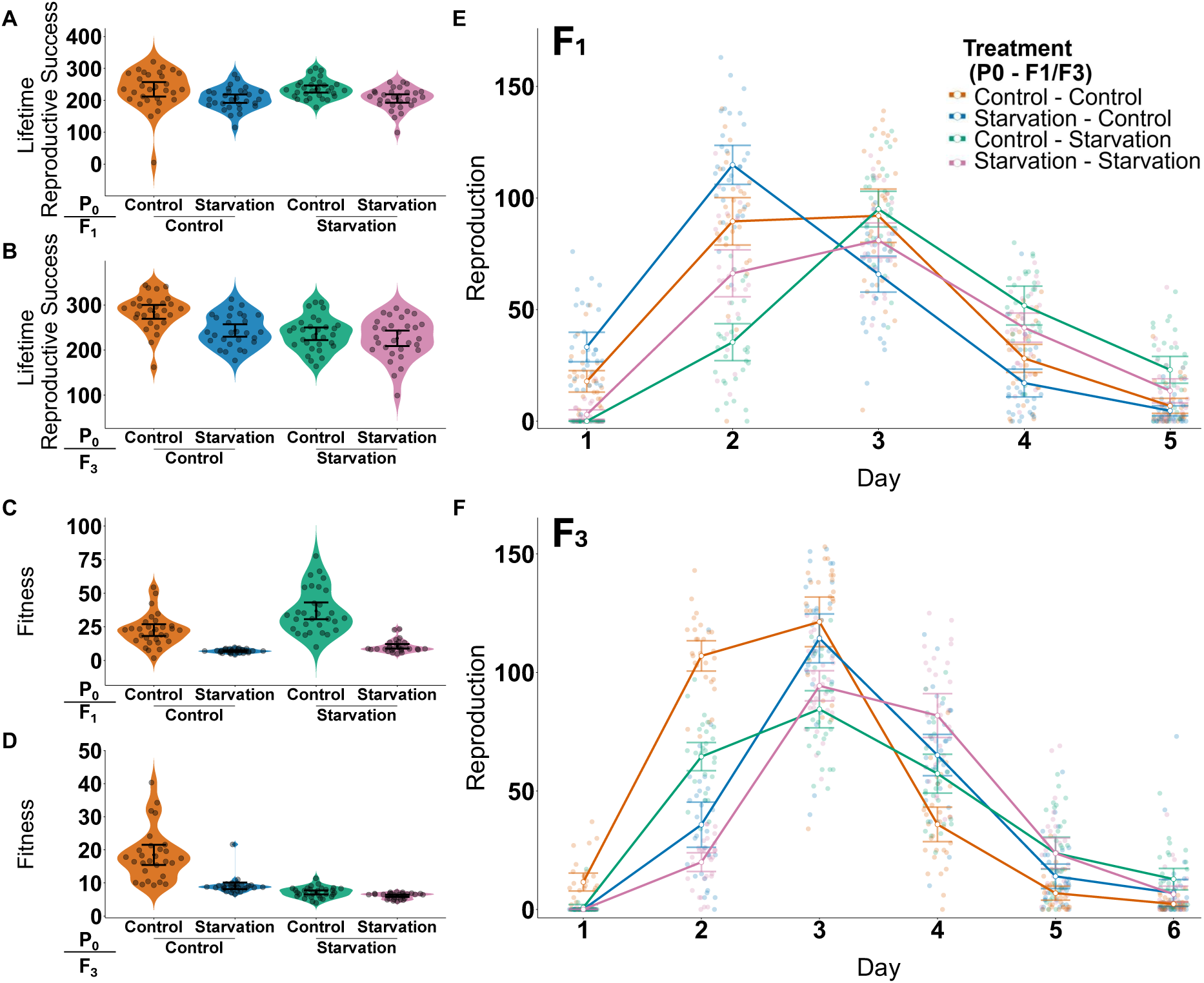
F1&F3 fitness assays: Violin plots show lifetime reproductive success (**A–B**) and rate sensitive fitness (λind) (**C–D**) of individuals from different treatment combinations across generations. (**E–F**) Daily reproduction across days of adulthood. Treatments represent combinations of parental P0 and F1 (**A,C,E**) or F3 (**B,D,F**). Plotted is daily reproduction, with line connecting the mean values for each treatment on each day. Each point represents one individual. All error bars represent 95% CIs.

#### Fitness

F_1_ offspring of starved parents showed improved fitness in several measures, regardless of the environment they themselves experienced. F_1_ individuals reared in control conditions had significantly higher lifetime reproductive success (LRS) than those reared in starvation conditions, irrespective of the parental treatment. *EMM wald z-test:* **F_1_*Control(P_0_Control) –* F_1_*Starvation(P_0_Control)****: z = 3.63, p = 0.0016;* **F_1_*Control(P_0_Control) –* F_1_*starvation(P_0_Starvation)****: z=3.21, p =0.0073;* **F_1_*Control(P_0_Control) –* F_1_*Starvation(P_0_Control)****: z = 3.36, p = 0.0043;* **F_1_*Control(P_0_Starvation) –* F_1_*Starvation(P_0_Starvation)****: z = 2.78, p = 0.027).* However, there was no difference in LRS in F_1_ individuals in control conditions regardless of parental treatments (*EMM wald z-test:* **F_1_Control(P_0_Control) – F_1_Control(P_0_Starvation)**: *z = 0.98, p =0.76, Figure 3A*). Similarly, there was no difference in LRS of individuals who underwent F_1_ larval starvation between parental treatments (*EMM wald z-test:* **F_1_*Starvation(P_0_Control) –* F_1_*Starvation(P_0_Control)****: z = -0.54, p = 0.95, Figure 3C*).

Rate-sensitive fitness was also significantly different between lineages (*Anova type III: χ2 =251.57, p < 0.001*) (*Figure 3B*). Similarly to LRS, individuals in F_1_ control conditions had higher λind than those who underwent F_1_ larval starvation regardless of parental treatment *(EMM wald z-test:* **F_1_*Control(P_0_Control) –* F_1_*Starvation(P_0_Control)****: z = 9.64, p < 0.001;* **F_1_*Control(P_0_Control) –* F_1_*Starvation(P_0_Starvation)****: z = 6.43, p < 0.001;* **F_1_*Control(P_0_Starvation) –* F_1_*Starvation(P_0_Control)****: z = 14.46, p < 0.001;* **F_1_*Control(P_0_Starvation) –* F_1_*Starvation(P_0_Starvation)****: z = 11.20, p < 0.001*). However, a comparison of worms in the same F_1_ environment shows that offspring of starved parents had higher λind than those from control parents (*EMM wald z-test:* **F_1_*Control(P_0_Control) –* F_1_*Control(P_0_Starvation)****: z = -4.79, p <0.001,* **F_1_*Starvation(P_0_Control) –* F_1_*Starvation(*F_1_*Starvation)****: z = -3.23, p = 0.0068*).

#### Survival

The parental diet of the F_1_ worms had a significant effect on offspring survival (*EMM wald z-test:* z = -2.86, p = 0.0042) (*Figure S2C*). However, the effect of offspring environment on survival was marginally non-significant (*EMM wald z-test:* z = 1.94, p = 0.052). Overall, in the F_1_ there was no significant interaction between offspring and parental treatment for survival (*EMM wald z-test:* z = 0.52, p = 0.61) (*Figure S2A*). Post-hoc analysis suggests that offspring in the control environment had increased survival if descended from starved parents compared to those descended from control parents (**F_1_*Control(P_0_Control) –* F_1_*Starvation(P_0_Control):*** *Log Odds = 0.67, 95%: 0.068, 1.27, p = 0.022*). However, in starved environments offspring from starved parents did not differ significantly in survival in comparison to those from control parents (**F_1_Starvation(P_0_Starvation) – F_1_Control(P_0_Starvation)**: *Log Odds = 0.51, 95%: -0.037, 1.06, p = 0.078*).

### F3

#### Age Specific Reproduction

Reproductive timing of great grand-offspring was significantly affected by ancestral treatment interactions, in terms of both the linear and quadratic components of age (*Figure 3F*) (*Anova type III: χ2 = 141.83, p < 0.001; χ2 = 106.95, p < 0.001*).

#### Fitness

Contrary to what was observed in the F_1_, in the F_3_, offspring of starved parents showed several decreased fitness estimates, regardless of the environment they themselves experienced. Individuals’ lifetime reproductive success was significantly altered by their lineage (*Anova type III: χ2 = 34.48, p <0.001*) (*Figure 3B*).

Specifically, regardless of ancestral treatment, individuals placed into control conditions from control great-grandparents had higher LRS than those who underwent F_3_ larval starvation (*EMM wald z-test: **F******_3_Control(P_0_Control) – F_3_Starvation(P_0_Control)****: z = 3.61, p = 0.0018, **F******_3_Control(P_0_Control) – F_3_Starvation(P_0_Starvation)****: z = 5.45, p < 0.001*). Furthermore, these individuals also had increased LRS compared to individuals placed in control environments from great-grandparents who underwent larval starvation (*EMM wald z-test: **F******_3_Control(P_0_Control) – F_3_Control(P_0_Starvation)****, z = 4.40, p < 0.001*). Conversely, the benefit to LRS when placed in a control environment was lost in individuals from larval starvation lineages, with no differences in LRS when comparison to individuals raised in larval starvation conditions *(EMM wald z-test: **F******_3_Control(P_0_Starvation) – F_3_Starvation(P_0_Control)****: z = -0.727, p = 0.89; **F******_3_Control(P_0_Starvation) – F_3_Starvation(P_0_Starvation)****: z = 1.11, p = 0.69*). Similarly, there was no effect of lineage on the LRS of individuals placed in larval starvation in the F_3_ (*EMM wald z-test: **F******_3_Starvation(P_0_Control) – F_3_Stavation(P_0_Starvation)****: z = 1.81, p = 0.27)*.

Great-grandparental lineage had a significant impact on rate-sensitive fitness (*Anova type III: χ2 =* 181.78, P < 0.001) (*Figure 3D*). Worms from a control great-grandparental lineage, placed in a control environment had increased *λind* when compared to individuals of any great-grandparental lineage that underwent larval starvation (*EMM wald z-test: **F******_3_*Control(P_0_Control) – F_3_Starvation(P_0_Control)**: z = 7.88, p < 0.001; **F_3_Control(P_0_Control) – F_3_Starvation(P_0_Starvation)**: z = 11.37, p < 0.001). This increase in *λind* was not observed in individuals from great-grandparental lineages which underwent larval starvation. (*EMM wald z-test: **F******_3_Control(P_0_Starvation) – F_3_Starvation(P_0_Control)****: z = -2.39, p = 0.080; **F******_3_Control(P_0_Starvation) – F3Starvation(P_0_Starvation)****: z = 1.42, p = 0.49*). However, when placed in the same environment, individuals from the control great-grandparental lineage had increased *λind* compared to those from larval starvation lineages (*EMM wald z-test: **F******_3_*Control(P_0_Control)** – **F_3_Control(P_0_Starvation)**: z= 10.21, p < 0.001; **F_3_Starvation(P_0_Control) – F_3_Starvation(P_0_Starvation)**: z = 3.76, p < 0.001).

#### Survival

The lineage of F_3_ worms had a significant effect on survival (*EMM wald z-test:* z = 2.08, p = 0.038) (Figure S2B) as well as the offspring treatment (*EMM wald z-test:* z =4.76, p < 0.001). The interaction between offspring and lineage treatment was significant for survival (*EMM wald z-test:* z = -2.62, 0.0087). Post-hoc analysis showed that F_3_ worms whose great-grandparents underwent larval starvation displayed no significant change in survival based on offspring environment (**F_3_*Starvation(P_0_Control) – F_3_Starvation(P_0_Starvation)****: log-odds = -1.33, 95%: - 1,02, 0.32, p =0.54*). In contrast, worms from well-fed great-grandparental lineages had decreased survival in starved conditions (**F_3_Control(P_0_Control) – F_3_Control(P_0_Starvation)** log-odds = -4.76, *95%: -1.99, 0.60* p= < 0.001) (*Figure S2D*).

#### Simulations of Transgenerational Trade-offs

We simulated a boom-and-bust population containing two strategies, one employing multigenerational trade-offs and one without. After 100 days both strategies persisted in the population with their frequencies oscillating (*Figure 4A*). However, individuals with multigenerational trade-offs consistently represented a higher average proportion of the population. After 1000 days, this advantage became more pronounced, with the multigenerational trade-off strategy almost reaching fixation (*Figure 4B*). These results show that multigenerational trade-offs can confer long-term adaptive benefits in transient boom-and-bust populations.

**Figure 4:**
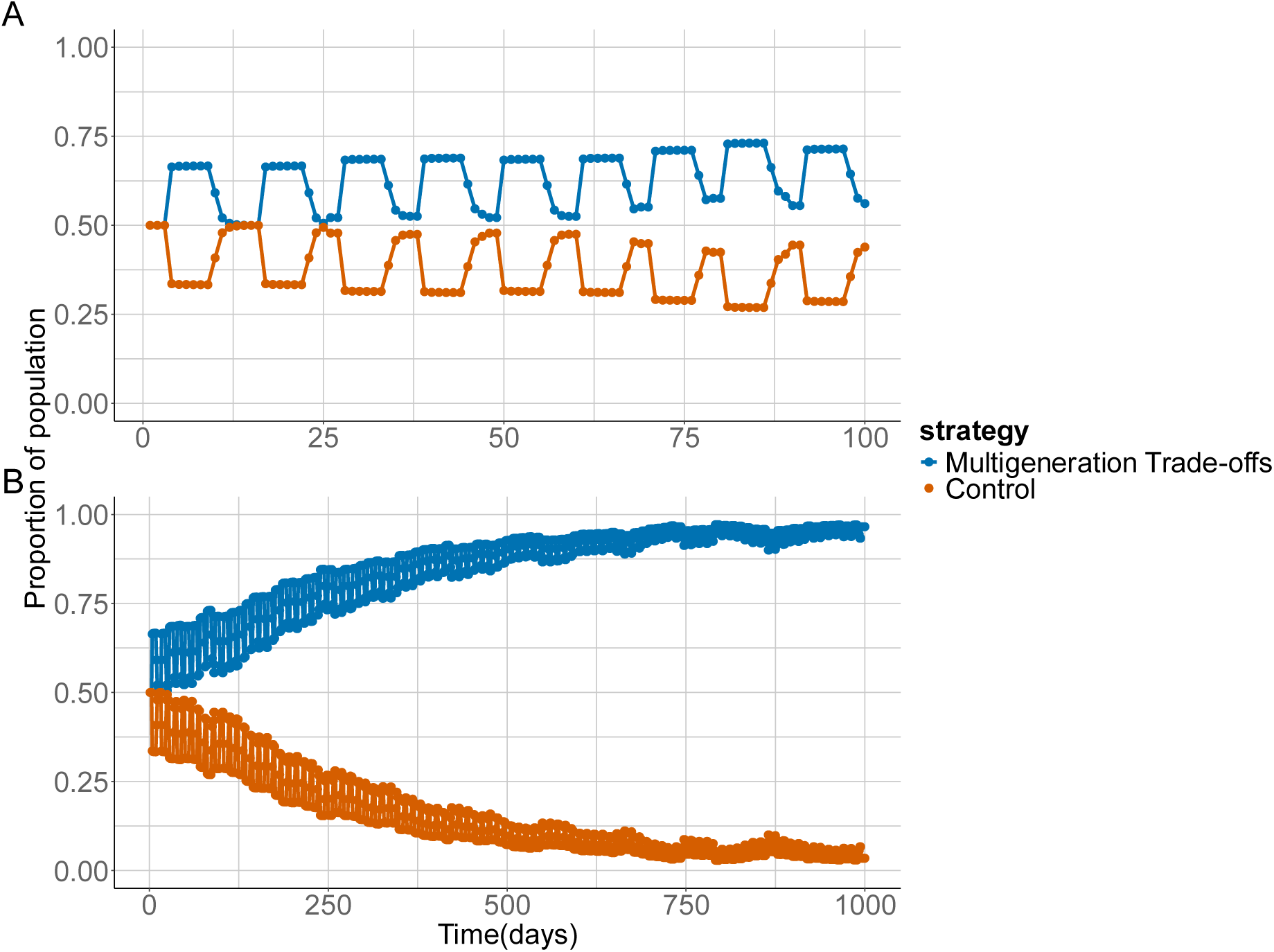
Evolutionary Simulations of multigenerational trade-offs. Plots show the output of evolutionary simulations of boom-and-bust populations after 100 days (**A**) and 1000 days (**B**). The simulations competed populations which utilized multigenerational trade-offs and those that did not.

## Discussion

Our results demonstrate that prolonged L1 arrest induced by starvation in *C. elegans* can induce adaptive intergenerational and maladaptive transgenerational effects. Across four complementary fitness read-outs, lifetime reproductive output, age-specific fecundity, rate-sensitive fitness (λ), and survival, we detected these effects in both matched and mis-matched environments, showing they are robust to the offspring’s rearing conditions. Our simulation model of boom-and-bust population dynamics further supports a novel evolutionary hypothesis, where short-term gains in the F_1_ generation are favoured even though they carry costs in the F_3_ generation, because of the net benefit to these alleles. In other words, selection on adaptive intergenerational plasticity can inadvertently cause maladaptive transgenerational outcomes that are tolerated because of the overall fitness benefit. While previous studies have found beneficial transgenerational effects, our results indicate that transgenerational effects need not be adaptive and underscore the importance of estimating fitness trade-offs across multiple generations and environments to fully resolve how organisms react to fluctuating environmental conditions. Importantly, we show that mal-adaptive transgenerational effects can evolve as a cost of selection on beneficial intergenerational effects.

Consistent with previous research, we show that starvation can induce adaptive intergenerational effects in *C. elegans*. Early-life starvation has been shown to cause reproductive defects in the parental generation^25^. However, offspring from starved worms are more resistant to starvation and less susceptible to germ line developmental abnormalities^27^. Importantly, F_1_ offspring of starved individuals reproduce more and earlier, regardless of environment, which, in a growing population, confers an evolutionary advantage ^28^. Our data suggests that the same environmental cue (here, starvation) that benefits F_1_ offspring may impose latent fitness costs two generations later. This finding cautions against assuming that intergenerational effects are uniformly beneficial even if they increase fitness in F_1_ generation. Instead, they can conceal delayed costs that only become visible when fitness is tracked beyond the first generation. Detecting such hidden costs will require the kind of multi-generation, multi-environment assays we deployed here. In this study, intergenerational benefits outweighed transgenerational costs, but this result may not be uniform across all taxa and environments. We suggest that more work is required to estimate the generality of such multigenerational trade-offs and evaluate their role in evolutionary processes. Transgenerational effects have been well documented, and *C. elegans* and provide a powerful tool for empirical tests of evolutionary hypotheses related to these effects. For instance, great-grand offspring from starved worms have been shown to have increased variability in stress-related phenotypes, which was hypothesized to be a form of diversifying bet-hedging_26,29_.

Additionally, great-grand offspring from worms that have undergone an extended dauer life-stage (an alternate dispersal life-stage) have increased starvation resistance, suggesting transgenerational anticipatory effects_20_. Theoretical work has shown that both bet-hedging and anticipatory effects are likely to evolve depending on the predictability of future environments_16,30_, and in some cases both strategies can be present simultaneously_31_. Both bet-hedging and anticipatory effects emphasize adaptive outcomes across generations. However, we show here that transgenerational effects can be maladaptive. Furthermore, our simulations suggest maladaptive transgenerational effects can be selected for if there are intergenerational benefits. The results outline a distinct evolutionary dynamic, that maladaptive transgenerational effects can emerge as a trade-off with intergenerational benefits. This contrasts with prior evolutionary models of transgenerational inheritance and provides a novel explanation for the existence of transgenerational effects.

Whilst experimental and theoretical work has identified adaptive intra-_22,32_ and intergenerational trade-offs_33,34_ longer term trade-offs have rarely been considered or documented. Our model shows that mal-adaptive transgenerational effects can evolve as a byproduct of selection on beneficial intergenerational effects when the transgenerational cost is mitigated by ecological dynamics. In *C.elegans*, for example, populations grow exponentially until they reach a local carrying capacity, at which point developing individuals enter an alternative dispersal life stage_35_. Such boom-and-bust population dynamics are common in nematodes and may mitigate the cost of mal-adaptive transgenerational effects facilitating their evolution, when there is a periodical resetting of environmental conditions. However, in a stable environment, transgenerational effects may have longer term fitness consequences, leading to different evolutionary outcomes (*Figure 5*). Testing for such context-dependent outcomes in taxa with contrasting population dynamics is therefore essential for determining how general multigenerational trade-offs are and which ecological settings favour their evolution.

**Figure 5:**
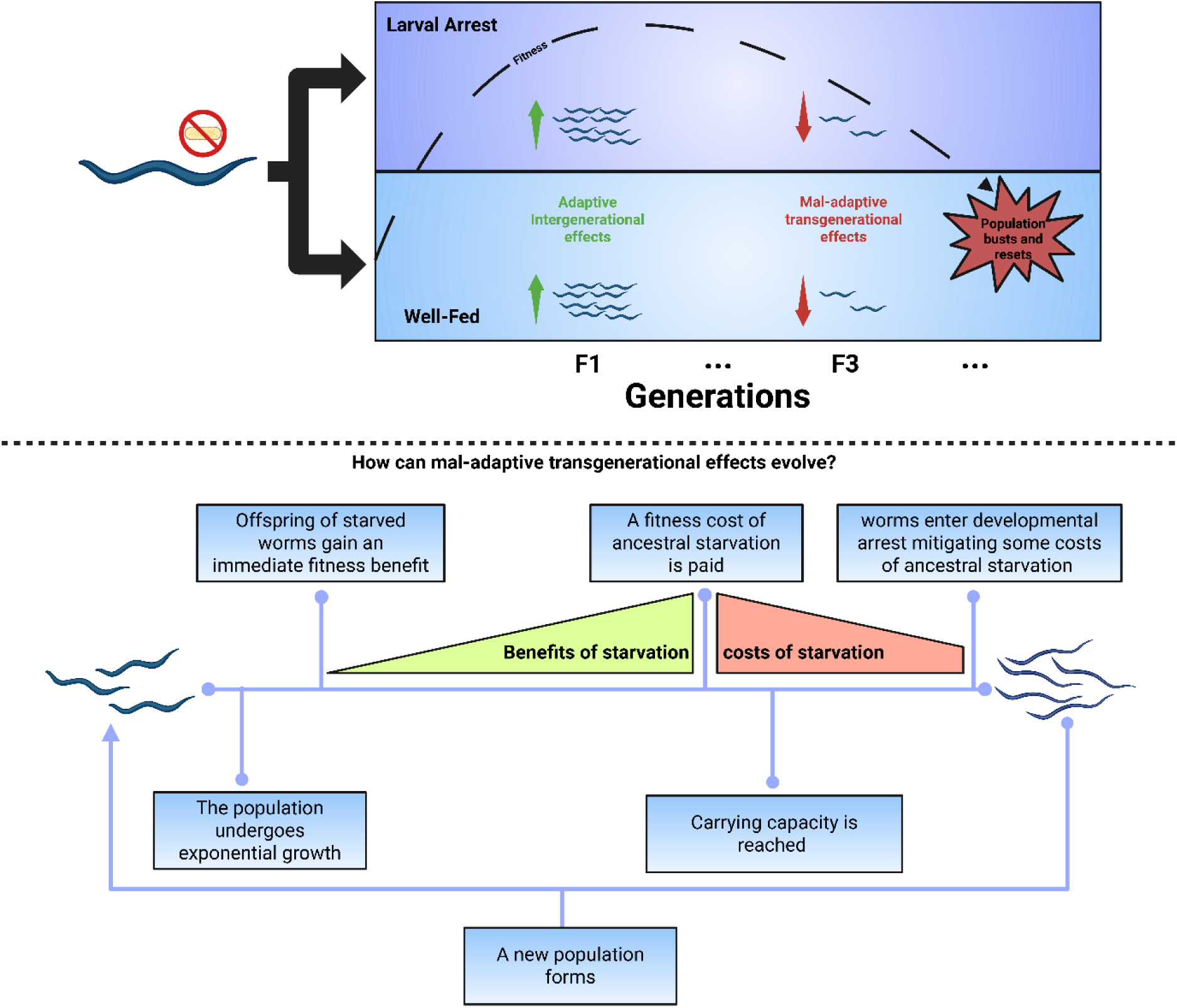
How can mal-adaptive transgenerational effects evolve? Illustration showing how larval starvation can generate both adaptive intergenerational and mal-adaptive transgenerational fitness effects. When larvae experience starvation and enter developmental arrest, their immediate offspring may exhibit adaptive intergenerational benefits. However, the cost of immediate beneficial effects is paid in a later generation. Worms enter developmental arrest due to the population reaching the carrying capacity, buffering some costs of ancestral starvation. Overall offspring receive more from the immediate benefits of larval starvation than future generations pay in eventual costs

We did not directly investigate the molecular mechanisms of transgenerational effects, but previous work has shown that larval starvation can induce the production of small RNAs that are inherited by great-grand offspring in *C.elegans*_36_. However, it is unlikely that multigenerational trade-offs are mediated through small RNAs alone. We suggest an alternative hypothesis, in which intergenerational effects arise from increased maternal provisioning, and transgenerational effects are due to a trade-off between transposable element (TE) activity and gene expression_37_. Previous work has shown that stressful environments can alter TE activity ^38^. Notably, the silencing of TEs can have bystander effects such that genes adjacent to silenced TEs show altered expression and can produce mal-adaptive phenotypes in later generations _37,39_. The “TE-mediated trade-off” model provides a framework to address unanswered questions about transgenerational effects. For instance, whilst our model still accommodates previous observations of sRNA inheritance, it can also explain mal-adaptive effects as a cost of silencing selfish genetic elements. However, although this model has some theoretical support_40_, future work is needed targeting small RNA pathways and TEs to determine the mechanisms which underlie long-term fitness costs of parental stress and shape multigenerational trade-offs.

Together, our results show that extended larval starvation in *C. elegans* results in beneficial intergenerational effects and detrimental transgenerational effects, regardless of environmental conditions experienced by the offspring. Eco-evolutionary simulation models show that such maladaptive F3 outcomes can arise as a by-product of selection for adaptive intergenerational plasticity, overturning the common assumption that transgenerational effects are inherently beneficial. These findings underscore the need to measure fitness across multiple generations and ecological contexts to fully understand adaptive plasticity. Little is known about multigenerational trade-offs in other taxa and closing this gap will require experiments that utilize multigenerational fitness assays in organisms with diverse life histories across the tree of life. Integrating multigenerational trade-offs with ecological and evolutionary perspectives will help us predict how populations navigate fluctuating environments across generations.

## Methods

### Strains and aintenance

*C. elegans* nematodes of the N2 Bristol strain were used in all assays. All nematodes were ordered from the *Caenorhabditis* Genetics Centre, which is funded by NIH Office of Research Infrastructure Programs (P40 OD010440). Populations were initiated from thawed N2 worms from -80°C and maintained for at least 3 generations before bleaching. When not undergoing treatments and during fitness assays, nematodes were maintained on NGM (Nematode Growth Medium) agar plates seeded with a lawn of *E. coli* OP50.1P (90mm for maintenance and 35mm for assays) and kept in climate chambers set to 20°C, 60% relative humidity and constant darkness. NGM agar contained antibiotics (100 ug ml-1 ampicillin) and a fungicide (100 ug ml-1 nystatin). See wormbook (http:www.wor book.org) for details on OP50 cultures, S-buffer and M9 recipes.

### Intergenerational and ransgenerational Fitness Assays

Our experimental design resulted in 10 different treatments, two parental treatments consisting of either a seven-day larval starvation or control conditions (fully fed), F1 and F3 offspring had either a matched or mismatched diet to their parents or great-grandparents, respectively. To control for different developmental times across treatments, worms are plated for reproduction regardless of life stage. Under control conditions, worms typically took 36-48hrs to reach Late-L4 after treatments, Therefore, this range was used to set-up reproduction. Fitness assays were set up by randomly picking individual late L4 worms onto individual 35mm NGM seeded plates. For the next 6-7 days every 24hr worms were transferred to new plates. Any eggs laid on the plates were allowed to develop for 2 days before being heat-shocked at 42°C for ∼3 hours and then counted. Worms were censored if they went missing, and worms that died before the end of reproduction (e.g. due to matricide) were scored as having 0 reproduction from that point.

Survival assays were conducted on 35mm seeded OP50.IP NGM plates. 10 worms were plated per plate, worms were transferred every 2 days, or 1 day during reproduction. Lifespan was checked every day; death was defined as the absence of worm movement after a light touch. A worm was censored if it died due to unnatural circumstances, such as matricides or going missing.

### Statistical Analysis and Simulations

All statistical analysis and simulations were conducted using R version 4.3.2_41_. For the P_0_ models, a fixed effect of treatment was fitted along with a random effect of ‘plate’ to account for potential pseudo-replication. For F_1_ and F_3_ models, Treatment consisted of the worm’s lineage resulting in a factor of 4 levels, ‘plate’ was fitted as a random variable.

We analysed 4 fitness estimates using generalised linear mixed effect models. The first of which, age specific reproduction, required further fixed effects of ‘Day’ along with a quadratic term of ‘Day’ and its interaction with ‘Treatment’. Furthermore, ‘Day’ was included as an additional random effect. Age-specific reproduction often shows significant dispersion and zero-inflation in *C. elegans*. Therefore, model diagnostics were run using the DHARMa package, to check for this_42_. Multiple error distributions were fitted using the GlmmTMB package (*see supplementary information*) and age-specific reproduction curves were visualised using GGplot2_43,44_.

The second estimate of fitness we analysed was lifetime reproductive success (*LRS*, total number of offspring produced per individual). Here, a basic generalized linear mixed model consisting of a negative binomial error structure and fixed effects of treatment, lineage (which treatment did their ancestor undergo) and a random effect of ‘Plate’ was used. We also fitted an alternative estimate of fitness, rate-sensitive fitness (*λind*), which can be estimated through extracting the dominant eigenvalue of individual structured Leslie matrices of life-history data _45_. The model was fitted, and data visualised in the same manner as LRS.

Our final fitness estimate was survival. Only natural deaths were included in the final analysis, matricides and lost worms were censored. Most data conformed to the assumptions of a Cox proportional hazard model, except data from the F_1_ worms. Therefore, we used an event history analysis for all data for consistency. Since the model requires sampling from every day, we added a random effect of ‘Worm ID’ to account for the repeated measures. The output from the model was then visualised through a forest plot and survival curve using ggplot2.

In all cases, Model selection was performed to identify the best zero-inflation and dispersion parameters for all response variables. AICs were then compared before the final model assumptions were checked using the DHARMa package. Finally, to determine the overall effect of lineage in the F_1_ and F_3_, pairwise tests were conducted using the emmeans function, to estimate marginal means_46_. If the model included an interaction term, then the emtrends function was used instead, which compares the estimated marginal slopes. Outputs from these functions were run through the pairs function for pairwise analyses.

Simulations were run using on RStudio_41_, based on collected data and grounded in existing knowledge of nematode ecology. The simulations explored the effects of competing two transgenerational strategies within a boom-and-bust nematode population. The first strategy simulated multigenerational trade-offs, which after starvation displayed F_1_ increases and F_3_ decreases in fitness with an intermediate point at F_2_. The other strategy exhibited no transgenerational effects, i.e. constant fitness. We assumed each strategy reacted the same way in the exposed (parental) generation, and the difference was in their intergeneration and transgenerational effects. The simulation input consists of ‘cohorts’ of worms containing information on the worms age, brood size, generations since starvation and transgenerational strategy. Global parameters determined the number of bacterial cells (food), age-specific reproduction/feeding and the relative fitness consequences of transgenerational effects (*Table S1*). The simulation was run through iterations, each representing one day in which the worms eat food, age, lay eggs and with new broods being added to the population size as a cohort, sharing parental transgenerational strategy, with the process being repeated until a set number of days was reached. When the population runs out of food a starvation event is triggered, and the worms enter the dispersal life stage. Worms in larval stages ‘disperse’ to find a new food source, the simulation now follows a new population formed of dispersed larvae. Simulation outputs were visualised using ggplot2.

## Acknowledgments

This work was supported by a grant from the Leverhulme Trust (Grant no. RPG-2023-068) “The mechanisms and adaptive value of transgenerational epigenetic effects”. Thanks goes to members of the Maklakov lab

