## Supplemental files for "Evolutionary trade-offs between intergenerational and transgenerational fitness effects"

**Methods**

**Bleaching and P0 Set-up**

Mixed stage worms were maintained on NGM agar plates for multiple generations. When a plates has a large population of eggs they are washed, with 3-4ml of M9 buffer, to remove unwanted adults. M9 was then pipetted off and discarded. Washing was repeated once more but the plate was agitated by lightly scraping a sterile pipette tip across the surface releasing stuck eggs. 3.5ml of M9 was then pipetted into sterile 15ml tubes. A bleach solution (made up of 1g NaOH, 5ml dH20 and 8ml NaClO 10%) was added to the tubes until the final volume was 5ml. Tubes were inverted to mix before being left for 10 minutes being inverted every 2 minutes to ensure movement of eggs. Tubes were then pelleted for 2 minutes at 1300 rpm and excess liquid was aspirated. The pellet was washed by adding M9 up to the final volume of 5ml before being inverted to resuspend and pelleted again for 2 minutes at 1300 rpm. This wash step was repeated three more times in which the last time the pellet was resuspended in 1ml of S-buffer.

10ul of the bleached eggs solution was pipetted onto unseeded 35mm NGM plates and left for 30 minutes to dry. These plates were then observed under a microscope allowing for estimates of eggs per ul. For L1 arrest treatments worms were placed into 5ml of virgin S-buffer at a 1 egg/ul. For control treatments, this same density was maintained but eggs were placed in a solution of 38 mg/ml of OP50.1P. All treatments were maintained at constant agitation using a Fisherbrand Platform Rotator set at speed 10 inside of a climate chamber set to normal laboratory conditions. After 7 days the L1 arrest worms were recovered on seeded 90mm NGM plates and maintained as normal. The control worms were recovered after the first larval stage (~12 hours) to synchronize their exit and development with larval starvation treatment worms. Control worms were recovered and maintained after treatment the same as L1 worms.

**Bacterial Food Cultures**

To feed worms in liquid culture, a 2X dilution master stock of *OP50.1P E. coli* (76 mg/ml) was created. In a 15ml tube, 5ml of antibiotic containing terrific broth (TB + 100ampicillin + streptomycin) and a single colony of OP50.1P was added and grown for 20hr at 37c without shaking. After 20 hrs the 15ml of grown OP50.1P was then added to a 1L bottle of antibiotic-resistant TB and grown for a further 24hr at 37c without tilting. The fully grown OP50 was separated into 45ml aliquots and pelleted at 4500rpm at 20c for 10 minutes. 90% of the supernatant was removed before resuspending the pellet in the remaining supernatant. The contents of the tubes were transferred into a tube of known mass before being re-pelleted at 4500rpm at 20c for 10 minutes. Spinning down the pellet, when necessary, all supernatant was removed. The mass of the tube with pellet was measured allowing for the mass of the pellet to be calculated. The dilution can be made by calculating the amount of antibiotic containing S-buffer needed to add to achieve target concentration. The bacterial stock was stored at 4c for up to a month.

**Setting-up lineages and lines**

P0 worms recovered from liquid cultures were then split into 3 distinct lines: (i) the P0 line contained worms which have either just gone through larval starvation or our control treatment. (ii) The F1 line, which is further split into two lineages corresponding with the P0 treatment the worms originate from. (iii) A F3 line, which follows the same logic of the F1 line.

Lines run independently of each other and, after splitting at P0, never coincide. As such, the only time the worms re-enter treatment will be either in the F1 or the F3 depending on their line. For the remainder of the experiment, all worms are maintained as normal on 90mm NGM agar plates. In the F1 & F3 lines the two lineages (Control & Larval Starvation) re-enter liquid culture treatment in match mismatched conditions, resulting in 4 treatments per line (Control Lineage - Control, Control lineage - Larval Starvation, Larval Starvation lineage – Larval Starvation, Larva starvation lineage – Control). F1 and F3 experimental treatments are conducted in the same way as previously described P0 treatments. Eggs entering control treatments remain in liquid culture until the L1 stage (~12 hours). Eggs entering larval-starvation treatments remain in treatment for 7 days.

**Figures**

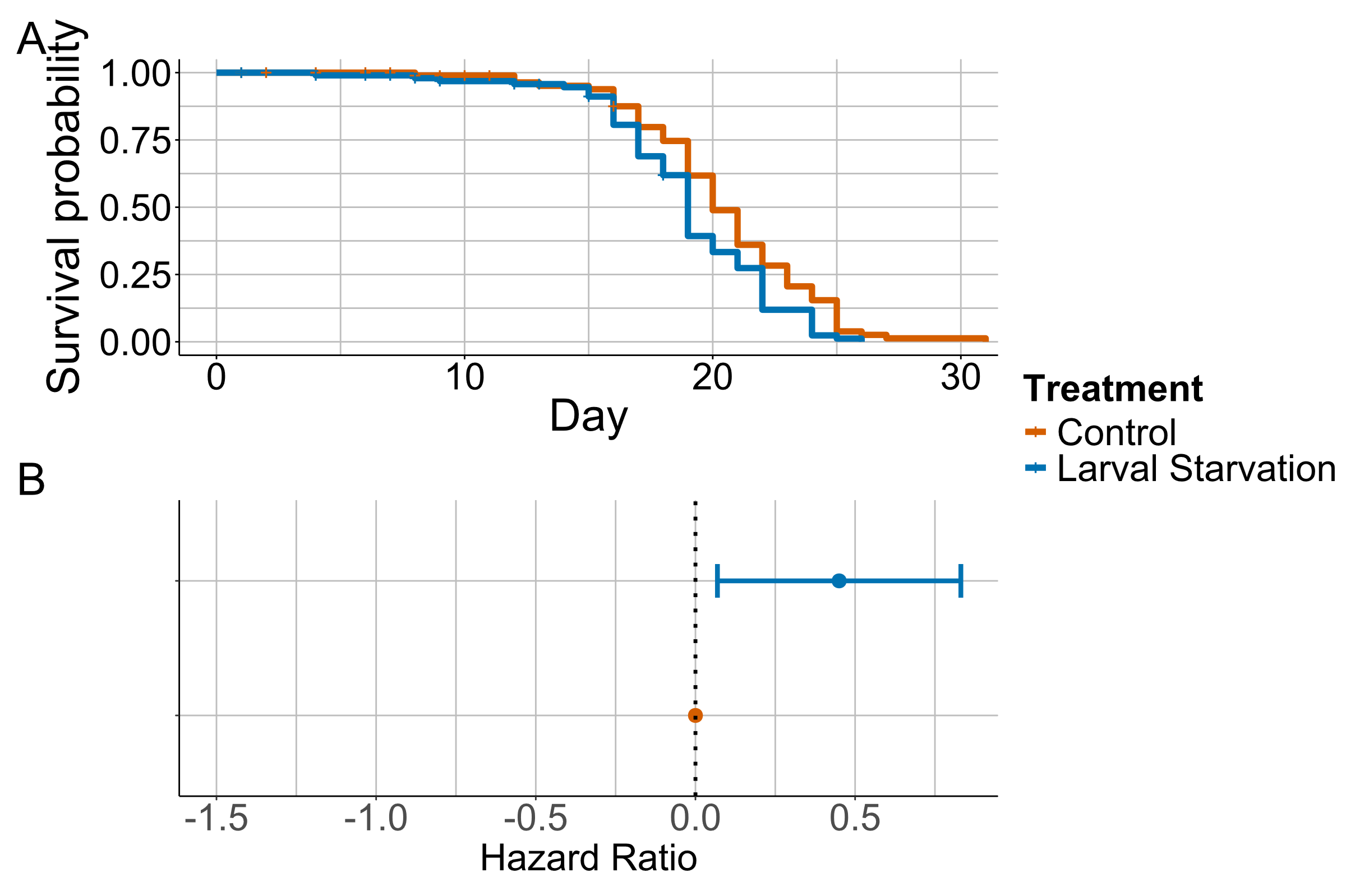

Figure S1: P0 worms developed in either control (ad-libitum) or larval starvation conditions. **A)** shows survival curves over the worms lifespan **B)** The hazard ratio. All error bars represent 95% CIs.

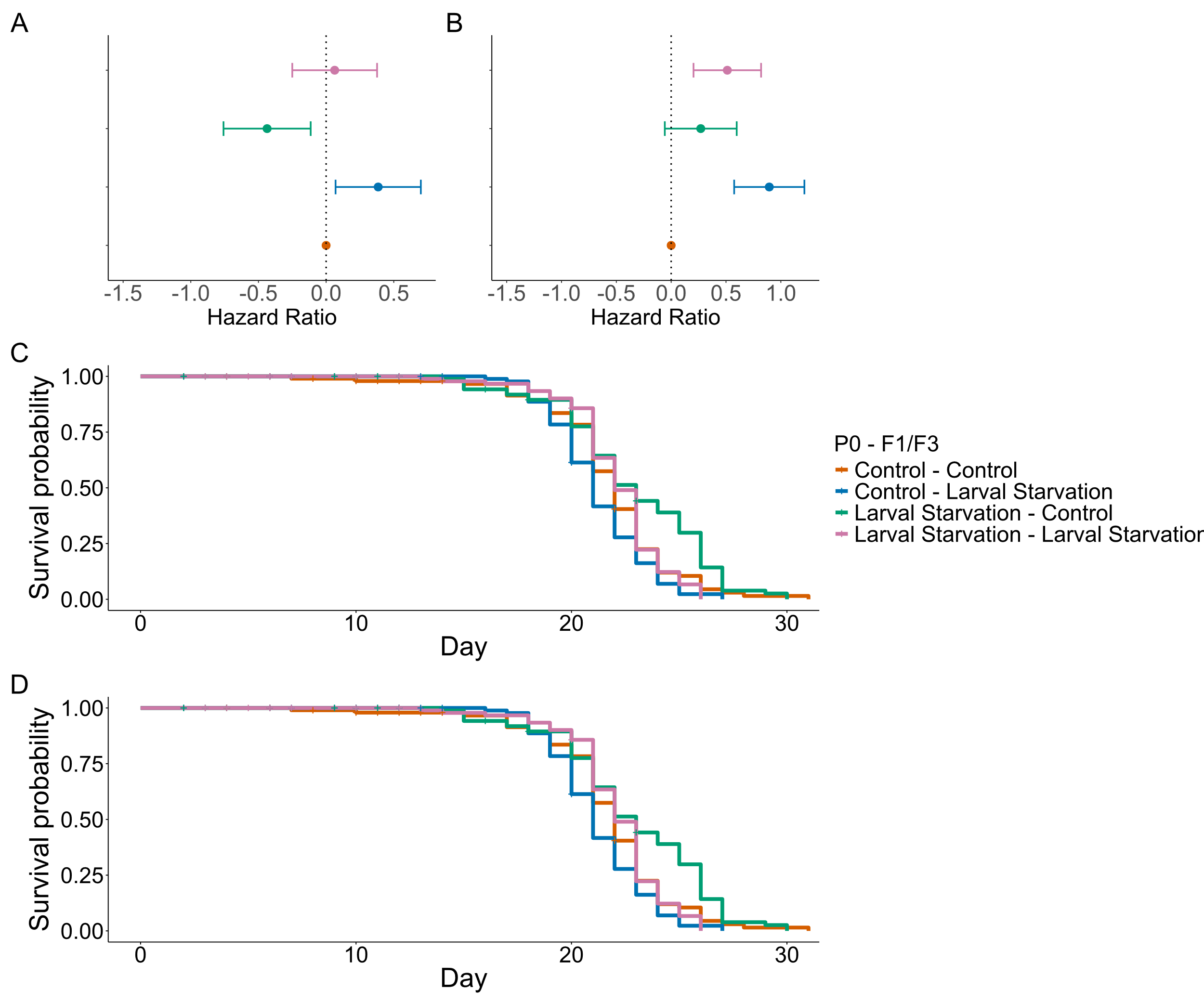

Figure S2: F1&F3 worms developed in either control (ad-libitum) or larval starvation conditions. **A)** The hazard ratio. **B)** shows survival curves over the worm’s lifespan. All error bars represent 95% CIs

Table S1: Example values of parameters used in the evolutionary simulation models

| **Parameter** | **Example** **Value** |
| --- | --- |
| Worm age | Variable |
| Cohort size | Variable |
| Generations since starvation | Variable |
| Transgenerational strategy | 1 for multigenerational trade-off, else 0 |
| Starting food | 1 x 10^8 – 10^10 |
| Reproduction | Varies based on age.  Day 1: 0  Day 2: 0  Day 3: 40  Day 4: 125  Day 5: 80  Day 6: 20  Day 7: 5  Day 8+: 0 |
| Feeding | Varies based on age.  Day 1: 1  Day 2: 1  Day 3: 2  Day 4: 2  Day 5: 4  Day 6: 4  Day 7: 4  Day 8+: 4 |

**Model Outputs**

**Age Specific Reproduction**

Table S2: P0 Age Specific Reproduction

Model formula: Reproduction ~ Treatment * poly(Day, 2) + (1 | Plate) + (1 | Day) | Dispersion: ~poly(Day, 2)

|  |  | Fixed Effects | | | | |  |
| --- | --- | --- | --- | --- | --- | --- | --- |
|  |  | Estimate | Std. Error | df | Statistic | p | Random Effects |
| **Predictors** | (Intercept) | 3.56 | 0.16 |  | 21.7 | <0.001 |  |
| **Predictors** | TreatmentLarval Starvation | -0.59 | 0.10 |  | -5.84 | <0.001 |  |
| **Predictors** | poly(Day, 2)1 | -13.18 | 2.87 |  | -4.59 | <0.001 |  |
| **Predictors** | poly(Day, 2)2 | -10.35 | 2.74 |  | -3.77 | <0.001 |  |
| **Predictors** | TreatmentLarval Starvation:poly(Day, 2)1 | 23.07 | 1.85 |  | 12.49 | <0.001 |  |
| **Predictors** | TreatmentLarval Starvation:poly(Day, 2)2 | -11.82 | 1.78 |  | -6.63 | <0.001 |  |
| **Random Effects** | N Plate |  |  |  |  |  | 58 |
| **Random Effects** | N Day |  |  |  |  |  | 58 |
| **Random Effects** | Observations |  |  |  |  |  | 58 |
| **ANOVA** | Treatment |  |  | 1 | Chisq = 33.39 | <0.001 |  |
| **ANOVA** | poly(Day, 2) |  |  | 2 | Chisq = 20.96 | <0.001 |  |
| **ANOVA** | Treatment:poly(Day, 2) |  |  | 2 | Chisq = 195.64 | <0.001 |  |

Table S3: F1 Age Specific Reproduction

Model formula: Reproduction ~ Treatment + poly(Day, 2) + (1 | Day) + (1 | Plate) + Treatment:poly(Day, 2) | Dispersion: ~(1 | Plate) + poly(Day, 2)

|  |  | Fixed Effects | | | | |  |
| --- | --- | --- | --- | --- | --- | --- | --- |
|  |  | Estimate | Std. Error | df | Statistic | p | Random Effects |
| **Predictors** | (Intercept) | 3.50 | 0.14 |  | 24.39 | <0.001 |  |
| **Predictors** | TreatmentC(L) | -0.11 | 0.08 |  | -1.41 | 0.157 |  |
| **Predictors** | TreatmentL(C) | -0.54 | 0.09 |  | -5.77 | <0.001 |  |
| **Predictors** | TreatmentL(L) | -0.22 | 0.08 |  | -2.76 | 0.006 |  |
| **Predictors** | poly(Day, 2)1 | -6.77 | 3.61 |  | -1.87 | 0.061 |  |
| **Predictors** | poly(Day, 2)2 | -20.28 | 3.54 |  | -5.73 | <0.001 |  |
| **Predictors** | TreatmentC(L):poly(Day, 2)1 | -12.72 | 2.06 |  | -6.18 | <0.001 |  |
| **Predictors** | TreatmentL(C):poly(Day, 2)1 | 32.33 | 2.49 |  | 13 | <0.001 |  |
| **Predictors** | TreatmentL(L):poly(Day, 2)1 | 14.43 | 2.20 |  | 6.55 | <0.001 |  |
| **Predictors** | TreatmentC(L):poly(Day, 2)2 | 4.04 | 1.92 |  | 2.1 | 0.035 |  |
| **Predictors** | TreatmentL(C):poly(Day, 2)2 | -11.91 | 2.32 |  | -5.13 | <0.001 |  |
| **Predictors** | TreatmentL(L):poly(Day, 2)2 | -4.53 | 2.07 |  | -2.19 | 0.028 |  |
| **Random Effects** | N Day |  |  |  |  |  | 5 |
| **Random Effects** | N Plate |  |  |  |  |  | 5 |
| **Random Effects** | Observations |  |  |  |  |  | 5 |
| **ANOVA** | Treatment |  |  | 3 | Chisq = 8.19 | 0.042 |  |
| **ANOVA** | poly(Day, 2) |  |  | 2 | Chisq = 28.65 | <0.001 |  |
| **ANOVA** | Treatment:poly(Day, 2) |  |  | 6 | Chisq = 395.43 | <0.001 |  |

Table S4: F3 Age Specific Reproduction

Model formula: Reproduction ~ Day + I(Day^2) + Treatment + (1 | Day) + Day:Treatment + I(Day^2):Treatment | Dispersion: ~poly(Day, 2) + Treatment + (1 | Plate) + (1 | Day) + poly(Day, 2):Treatment

|  |  | Fixed Effects | | | | |  |
| --- | --- | --- | --- | --- | --- | --- | --- |
|  |  | Estimate | Std. Error | df | Statistic | p | Random Effects |
| **Predictors** | (Intercept) | 0.81 | 0.79 |  | 1.02 | 0.307 |  |
| **Predictors** | Day | 2.56 | 0.51 |  | 4.99 | <0.001 |  |
| **Predictors** | I(Day^2) | -0.44 | 0.07 |  | -6.11 | <0.001 |  |
| **Predictors** | TreatmentC(L) | -4.21 | 0.61 |  | -6.9 | <0.001 |  |
| **Predictors** | TreatmentL(C) | -0.26 | 0.36 |  | -0.72 | 0.473 |  |
| **Predictors** | TreatmentL(L) | -5.74 | 0.47 |  | -12.19 | <0.001 |  |
| **Predictors** | Day:TreatmentC(L) | 2.00 | 0.34 |  | 5.8 | <0.001 |  |
| **Predictors** | Day:TreatmentL(C) | -0.44 | 0.23 |  | -1.93 | 0.054 |  |
| **Predictors** | Day:TreatmentL(L) | 2.54 | 0.27 |  | 9.29 | <0.001 |  |
| **Predictors** | I(Day^2):TreatmentC(L) | -0.21 | 0.05 |  | -4.32 | <0.001 |  |
| **Predictors** | I(Day^2):TreatmentL(C) | 0.14 | 0.03 |  | 4.14 | <0.001 |  |
| **Predictors** | I(Day^2):TreatmentL(L) | -0.23 | 0.04 |  | -6.1 | <0.001 |  |
| **Random Effects** | N Day |  |  |  |  |  | 6 |
| **Random Effects** | Observations |  |  |  |  |  | 6 |
| **ANOVA** | Day |  |  | 1 | Chisq = 30.58 | <0.001 |  |
| **ANOVA** | I(Day^2) |  |  | 1 | Chisq = 40.96 | <0.001 |  |
| **ANOVA** | Treatment |  |  | 3 | Chisq = 97.34 | <0.001 |  |
| **ANOVA** | Day:Treatment |  |  | 3 | Chisq = 141.83 | <0.001 |  |
| **ANOVA** | I(Day^2):Treatment |  |  | 3 | Chisq = 106.96 | <0.001 |  |

**Lifetime Reproductive success**
Table S5: P0 Lifetime Reproductive success

Model formula: Reproduction ~ Treatment + (1 | Plate) | Dispersion: ~Treatment + 1 | Zero-inflation: ~1

|  |  | Fixed Effects | | | | |  |
| --- | --- | --- | --- | --- | --- | --- | --- |
|  |  | Estimate | Std. Error | df | Statistic | p | Random Effects |
| **Predictors** | (Intercept) | 5.60 | 0.02 |  | 254.41 | <0.001 |  |
| **Predictors** | TreatmentLarval Starvation | -0.29 | 0.07 |  | -4.25 | <0.001 |  |
| **Predictors** | (Intercept) | -21.67 | 6682.00 |  | 0 | 0.997 |  |
| **Random Effects** | N Plate |  |  |  |  |  | 58 |
| **Random Effects** | Observations |  |  |  |  |  | 58 |
| **ANOVA** | Treatment |  |  | 1 | Chisq = 18.04 | <0.001 |  |

Table S6: F1 Lifetime Reproductive Success

Model formula: Reproduction ~ Treatment + (1 | Plate) | Dispersion: ~Treatment + (1 | Plate)

|  |  | Fixed Effects | | | | |  |
| --- | --- | --- | --- | --- | --- | --- | --- |
|  |  | Estimate | Std. Error | df | Statistic | p | Random Effects |
| **Predictors** | (Intercept) | 5.50 | 0.04 |  | 138.75 | <0.001 |  |
| **Predictors** | TreatmentC(L) | -0.05 | 0.05 |  | -0.98 | 0.329 |  |
| **Predictors** | TreatmentL(C) | -0.18 | 0.05 |  | -3.63 | <0.001 |  |
| **Predictors** | TreatmentL(L) | -0.15 | 0.05 |  | -3.21 | 0.001 |  |
| **Random Effects** | N Plate |  |  |  |  |  | 119 |
| **Random Effects** | Observations |  |  |  |  |  | 119 |
| **ANOVA** | Treatment |  |  | 3 | Chisq = 21.55 | <0.001 |  |

Table S7: F3 Lifetime Reproductive Success

Model formula: Reproduction ~ Treatment + (1 | Plate)

|  |  | Fixed Effects | | | | |  |
| --- | --- | --- | --- | --- | --- | --- | --- |
|  |  | Estimate | Std. Error | df | Statistic | p | Random Effects |
| **Predictors** | (Intercept) | 5.65 | 0.03 |  | 189.49 | <0.001 |  |
| **Predictors** | TreatmentC(L) | -0.19 | 0.04 |  | -4.4 | <0.001 |  |
| **Predictors** | TreatmentL(C) | -0.16 | 0.04 |  | -3.61 | <0.001 |  |
| **Predictors** | TreatmentL(L) | -0.24 | 0.04 |  | -5.47 | <0.001 |  |
| **Random Effects** | N Plate |  |  |  |  |  | 116 |
| **Random Effects** | Observations |  |  |  |  |  | 116 |
| **ANOVA** | Treatment |  |  | 3 | Chisq = 34.48 | <0.001 |  |

**Rate Sensitive Fitness**

Table S8: P0 Rate Sensitive Fitness

Model formula: dominant_eigenvalues ~ Treatment + (1 | Plate)

|  |  | Fixed Effects | | | | |  |
| --- | --- | --- | --- | --- | --- | --- | --- |
|  |  | Estimate | Std. Error | df | Statistic | p | Random Effects |
| **Predictors** | (Intercept) | 3.76 | 0.05 |  | 78.13 | <0.001 |  |
| **Predictors** | TreatmentLarval Starvation | -1.68 | 0.07 |  | -24.13 | <0.001 |  |
| **Random Effects** | N Plate |  |  |  |  |  | 58 |
| **Random Effects** | Observations |  |  |  |  |  | 58 |
| **ANOVA** | Treatment |  |  | 1 | Chisq = 582.12 | <0.001 |  |

Table S9: F1 Rate Sensitive Fitness

Model formula: dominant_eigenvalues ~ Treatment + (1 | Plate)

|  |  | Fixed Effects | | | | |  |
| --- | --- | --- | --- | --- | --- | --- | --- |
|  |  | Estimate | Std. Error | df | Statistic | p | Random Effects |
| **Predictors** | (Intercept) | 3.00 | 0.08 |  | 38.66 | <0.001 |  |
| **Predictors** | TreatmentC(L) | 0.52 | 0.11 |  | 4.79 | <0.001 |  |
| **Predictors** | TreatmentL(C) | -1.08 | 0.11 |  | -9.64 | <0.001 |  |
| **Predictors** | TreatmentL(L) | -0.72 | 0.11 |  | -6.43 | <0.001 |  |
| **Random Effects** | N Plate |  |  |  |  |  | 119 |
| **Random Effects** | Observations |  |  |  |  |  | 119 |
| **ANOVA** | Treatment |  |  | 3 | Chisq = 251.57 | <0.001 |  |

Table S10: F3 Rate Sensitive Fitness

Model formula: dominant_eigenvalues ~ Treatment + (1 | Plate)

|  |  | Fixed Effects | | | | |  |
| --- | --- | --- | --- | --- | --- | --- | --- |
|  |  | Estimate | Std. Error | df | Statistic | p | Random Effects |
| **Predictors** | (Intercept) | 2.90 | 0.05 |  | 52.76 | <0.001 |  |
| **Predictors** | TreatmentC(L) | -0.94 | 0.09 |  | -10.2 | <0.001 |  |
| **Predictors** | TreatmentL(C) | -0.70 | 0.09 |  | -7.88 | <0.001 |  |
| **Predictors** | TreatmentL(L) | -1.10 | 0.10 |  | -11.37 | <0.001 |  |
| **Random Effects** | N Plate |  |  |  |  |  | 116 |
| **Random Effects** | Observations |  |  |  |  |  | 116 |
| **ANOVA** | Treatment |  |  | 3 | Chisq = 181.78 | <0.001 |  |

**Survival**

Table S21: P0 survival

Model formula: status ~ Treatment + Age + (1 | Plate/Worm) + (1 | Age)

|  |  | Fixed Effects | | | | |  |
| --- | --- | --- | --- | --- | --- | --- | --- |
|  |  | Estimate | Std. Error | df | Statistic | p | Random Effects |
| **Predictors** | (Intercept) | -9.76 | 1.04 |  | -9.42 | <0.001 |  |
| **Predictors** | TreatmentL1 | 0.61 | 0.26 |  | 2.31 | 0.021 |  |
| **Predictors** | Age | 0.41 | 0.06 |  | 7.16 | <0.001 |  |
| **Random Effects** | N :WormPlate |  |  |  |  |  | 0 |
| **Random Effects** | N Plate |  |  |  |  |  | 0 |
| **Random Effects** | N Age |  |  |  |  |  | 0 |
| **Random Effects** | Observations |  |  |  |  |  | 0 |
| **ANOVA** | Treatment |  |  | 1 | Chisq = 5.33 | 0.021 |  |
| **ANOVA** | Age |  |  | 1 | Chisq = 51.24 | <0.001 |  |

Table S32: F1 survival

Model formula: status ~ Treatment * Parental.Treatment + Age + (1 | Plate/Worm) + (1 | Age)

|  |  | Fixed Effects | | | | |  |
| --- | --- | --- | --- | --- | --- | --- | --- |
|  |  | Estimate | Std. Error | df | Statistic | p | Random Effects |
| **Predictors** | (Intercept) | -12.13 | 1.06 |  | -11.48 | <0.001 |  |
| **Predictors** | TreatmentL1 | 0.43 | 0.22 |  | 1.94 | 0.052 |  |
| **Predictors** | Parental.TreatmentL1 | -0.67 | 0.23 |  | -2.86 | 0.004 |  |
| **Predictors** | Age | 0.50 | 0.05 |  | 9.25 | <0.001 |  |
| **Predictors** | TreatmentL1:Parental.TreatmentL1 | 0.16 | 0.31 |  | 0.52 | 0.606 |  |
| **Random Effects** | N :WormPlate |  |  |  |  |  | 0 |
| **Random Effects** | N Plate |  |  |  |  |  | 0 |
| **Random Effects** | N Age |  |  |  |  |  | 0 |
| **Random Effects** | Observations |  |  |  |  |  | 0 |
| **ANOVA** | Treatment |  |  | 1 | Chisq = 10.69 | 0.001 |  |
| **ANOVA** | Parental.Treatment |  |  | 1 | Chisq = 12.74 | <0.001 |  |
| **ANOVA** | Age |  |  | 1 | Chisq = 85.58 | <0.001 |  |
| **ANOVA** | Treatment:Parental.Treatment |  |  | 1 | Chisq = 0.27 | 0.606 |  |

Table S43: F3 survival

Model formula: status ~ Parental.Treatment * Treatment + Age + (1 | Plate/Worm) + (1 | Age)

|  |  | Fixed Effects | | | | |  |
| --- | --- | --- | --- | --- | --- | --- | --- |
|  |  | Estimate | Std. Error | df | Statistic | p | Random Effects |
| **Predictors** | (Intercept) | -12.20 | 1.22 |  | -10.02 | <0.001 |  |
| **Predictors** | Parental.TreatmentL1 | 0.54 | 0.26 |  | 2.08 | 0.038 |  |
| **Predictors** | TreatmentL1 | 1.29 | 0.27 |  | 4.76 | <0.001 |  |
| **Predictors** | Age | 0.50 | 0.06 |  | 8.05 | <0.001 |  |
| **Predictors** | Parental.TreatmentL1:TreatmentL1 | -0.94 | 0.36 |  | -2.62 | 0.009 |  |
| **Random Effects** | N :WormPlate |  |  |  |  |  | 0 |
| **Random Effects** | N Plate |  |  |  |  |  | 0 |
| **Random Effects** | N Age |  |  |  |  |  | 0 |
| **Random Effects** | Observations |  |  |  |  |  | 0 |
| **ANOVA** | Parental.Treatment |  |  | 1 | Chisq = 0.05 | 0.830 |  |
| **ANOVA** | Treatment |  |  | 1 | Chisq = 16.61 | <0.001 |  |
| **ANOVA** | Age |  |  | 1 | Chisq = 64.87 | <0.001 |  |
| **ANOVA** | Parental.Treatment:Treatment |  |  | 1 | Chisq = 6.87 | 0.009 |  |
